# Phylogenetic diversity and species diversity are decoupled under experimental warming and cooling in Rocky Mountain plant communities

**DOI:** 10.1101/2025.06.11.659139

**Authors:** Leah N Veldhuisen, Lorah Seltzer, Jocelyn Navarro, Brian J. Enquist, Katrina M Dlugosch

## Abstract

**Aim:** Nearly 8% of species could go extinct from climate change; many organisms are already experiencing declines in abundance, local extinction, and range shifts. How such changes impact community diversity is an open question in most systems. Whether changes in phylogenetic diversity parallel those in traditional diversity metrics is also often unknown. We used experimentally transplanted plant communities to ask how different aspects of community diversity change with environmental factors across elevation, and whether phylogenetic relationships predict individual species’ responses to change.

**Location:** We experimentally transplanted subalpine plant communities down and upslope across 400 m of elevation at the Rocky Mountain Biological Laboratory, Gothic, CO, USA, to simulate climate warming and cooling.

**Methods:** We identified how experimental warming and cooling impacted community diversity by testing for differences in species richness, Shannon diversity, and phylogenetic diversity among transplant treatments. We tested for phylogenetic signal in each species’ change in percent cover among treatments. Finally, we assessed if aspects of species’ rarity (and thus their putative extinction risk) predicted post-transplant change independently or in addition to their phylogenetic relationship within the community.

**Results:** We found that species richness and Shannon diversity decreased in cooled treatments and increased in some warmed treatments. In contrast, phylogenetic diversity increased in the cooled treatment and did not change in the warmed treatments. Individual species’ changes in response to warming and cooling were not correlated with phylogeny or aspects of rarity.

**Main conclusions:** Our results suggest that species losses in cooled treatments are phylogenetically dispersed, increasing phylogenetic diversity, even as richness and Shannon diversity decline. Increasing richness and Shannon diversity in warmed treatments suggests that new species from across the phylogeny can colonize after transplantation, leading to stability in phylogenetic diversity under warming at this time scale, with further change likely as extinction debts are paid.

## Introduction

Globally, it has been predicted that nearly 8% of species could be driven extinct by climate change(Urban, 2024). Populations of many organisms are already experiencing significant changes. For example, in Thoreau’s Woods in the northeastern United States, researchers have documented declines in abundance, increased local extinction, and advancing phenology in the plant community (Willis et al., 2008). Similarly, over half of mammal species studied in the southern Rocky Mountains have already shifted their elevational ranges upward (McCain et al., 2021), and other taxonomic groups are experiencing similar range shifts in response to climate warming (Chen et al., 2011; Zorio et al., 2016). While it is well established that community composition is changing and populations are declining (Parmesan & Yohe, 2003; Román-Palacios & Wiens, 2020), the details of which species are lost and the impact on community composition remains poorly understood in most systems (Chichorro et al., 2019; Hordijk et al., 2024; Purvis et al., 2000; Staude, Waller, et al., 2020; Torres-Romero et al., 2023; Van der Colff et al., 2023; Wilder et al., 2023).

Phylogenetic diversity, the variation in evolutionary relationships present in a community, may be able to inform and predict the impacts of changes in community composition. Phylogenetic diversity is positively associated with ecosystem stability and productivity (Cadotte, 2013; Cadotte et al., 2008, 2012; Davies et al., 2016), and is more positively correlated with biomass than with species richness or functional diversity (Cadotte et al., 2008). It can also increase resistance to invasion if functional diversity is also high (Galland et al., 2019). More broadly, phylogenetic diversity can capture variation among species that is missed when only studying one or a few traits, which could potentially more accurately reflect niche space (Cadotte et al., 2009; E-Vojtkó et al., 2023; Huang et al., 2020). Finally, phylogenetic diversity, is predicted to decline with phylogenetically clustered extinction and extinction cascades (Eiserhardt et al., 2015; González-Orozco et al., 2016; Li et al., 2019; Thébault et al., 2007).

Despite the role of phylogenetic diversity in ecosystem functioning and its conservation importance (Cadotte, 2013; Cadotte et al., 2008, 2012; Davies et al., 2016; Molina-Venegas et al., 2020; Owen et al., 2019), previous work has shown that it does not respond consistently to global change (Daru et al., 2021; Padullés Cubino et al., 2024; Winter et al., 2009), and cannot be assumed to correlate with functional diversity (E-Vojtkó et al., 2023; Gerhold et al., 2015; Hähn et al., 2024). A recent study of European understory plants found that phylogenetic diversity did not change over a 40 year period, and that lost and gained species were randomly dispersed across the phylogeny (Padullés Cubino et al., 2024). In contrast, Li and colleagues (2019) found that grassland communities lost phylogenetic diversity over a 19-year period due to decreased winter precipitation, but only at the local scale. Both extinctions and invasions contribute to changes in community diversity, as invaders and emigrants can increase or decrease phylogenetic diversity, depending on their relatedness to the original community (Bennett et al., 2014; Daru et al., 2021; Winter et al., 2009). Other previous work shows that functional and phylogenetic diversity can respond in opposite directions to the same environmental variable (De Pauw et al., 2021), while Li et al. (2019) and Miller et al. (2019) found that phylogenetic and functional diversity both declined in their common study system over the same time frame. Given the ecosystem benefits of phylogenetic diversity and its independence of other diversity metrics, it is critical to better understand how it may respond to climate change (White et al., 2023).

Species’ occurrences determine community composition, so understanding which species survive or die can clarify and predict community-level change. A species’ evolutionary history may influence its response to climate change (Dinnage et al., 2020; Molina-Venegas et al., 2020; Padullés Cubino et al., 2024; Vamosi & Wilson, 2008). Evolutionarily distinct species are often the most threatened (Greenberg et al., 2018; Johnson et al., 2002; Verde Arregoitia et al., 2013), and if species-poor clades have higher extinction risk, extinction will threaten the entire clade (Vamosi & Wilson, 2008). Certain traits may be extinction-prone and phylogenetically conserved, such that closely related species with similar traits will face a similar risk of extinction (Eiserhardt et al., 2015; Li et al., 2019). For example, in globally-distributed northern temperate trees, cold tolerance is associated with extinction risk and is phylogenetically conserved, leading to phylogenetically clustered extinction (Eiserhardt et al., 2015).

Extinction is often associated with a species’ rarity (Harnik et al., 2012; Manes et al., 2021; Purvis et al., 2000), but the relationship between rarity and phylogenetic composition of communities is also unclear (Herzog & Latvis, 2022; Veldhuisen et al., 2024). Both low abundance and small range size have been found to be closely associated with extinction(Harnik et al., 2012; Lucas et al., 2019; Staude, Navarro, et al., 2020; Staude, Waller, et al., 2020). Species with restricted ranges typically have high habitat specificity and limited dispersal ability, increasing the threat of habitat fragmentation and increased habitat heterogeneity (Alzate & Onstein, 2022; Chichorro et al., 2019; Harnik et al., 2012; Lucas et al., 2019; Purvis et al., 2000; Staude, Navarro, et al., 2020).

Here, we use a set of experimentally transplanted plant communities to simulate global change and investigate its consequences for different aspects of community diversity. We leverage transplanted turfs at the Rocky Mountain Biological Laboratory (Gothic, CO, USA) that are part of a worldwide network of similar transplants (Bektaş et al., 2023; Block et al., 2022; Haider et al., 2022; Henn et al., 2018; Lynn et al., 2023; Nomoto & Alexander, 2021; Vandvik et al., 2020). Transplant experiments are powerful tools for testing the relationships between communities and their environment (Bektaş et al., 2023; Block et al., 2022; Haider et al., 2022; Henn et al., 2018; Lynn et al., 2023; Nomoto & Alexander, 2021; Vandvik et al., 2020). The elevational gradient captures temperature, as well as other variables that change over elevation and are changing with climate change (Aldridge et al., 2011; Bryant et al., 2008; Prather et al., 2023), such as precipitation, soil moisture, soil temperature and snow melt date. These turfs are especially well-suited to address questions about species loss and phylogenetic diversity, as mountain regions are predicted to face some of the highest risk of extinction from climate change (Manes et al., 2021; Urban, 2024). Specifically, we test how transplants downslope (warming) and transplants upslope (cooling) impact community diversity by quantifying species richness, Shannon diversity, and metrics of phylogenetic diversity. We then quantify each species’ change in percent cover in response to transplanting, and test for phylogenetic signal in this change. We also assess whether aspects of rarity (and therefore putative extinction risk), including initial abundance or range size, can predict species’ post-transplant change, either as part of or independently from their relationship to phylogenetic diversity. Specifically we predict:

1) *Experimental warming will result in the decline of community richness and Shannon diversity, and cooling will cause little change.* Warming is leading to the decline of subalpine species, and this effect is expected to be particularly severe for high elevation communities that are adapted to the coolest temperatures (Manes et al., 2021; Urban, 2024). In contrast, experimental cooling is expected to lead to little change or even increases in richness and Shannon diversity as species persist or recruit to turfs under more favorable environments (Lynn et al., 2021, 2023). The effects of warming are already evident in our study communities (CaraDonna et al., 2014; Dunne et al., 2003; Panetta et al., 2018; Rivest et al., 2023; Zorio et al., 2016) and experimental cooling may restore favorable conditions.
2) *Experimental warming will result in the decline of phylogenetic diversity, and cooling will again cause little change.* If related species are lost together, then we predict that phylogenetic metrics will decline with warming and that abundance changes of individual species will have a phylogenetic signal. Previous work in this system found that species from certain clades disproportionately flower at warmer, drier points in the growing season (Veldhuisen et al., 2023), suggesting that traits shaping success or failure in warmer environments will be phylogenetically clustered. Alternatively, if phylogeny does not predict species decline in the transplants, then species losses will be dispersed across the phylogeny and there will be no phylogenetic signal in declines. Losses of species from across the phylogeny could result in a counterintuitive increase in some metrics of phylogenetic diversity in warmed treatments, as phylogenetic distances between remaining taxa increase. In cooled treatments, again we predict little change in phylogenetic diversity, as species are retained in more favorable conditions.
3) *Aspects of rarity will predict species declines under experimental warming.* We predict that aspects of rarity often associated with extinction, including both lower abundance and smaller range size (Manes et al., 2021; Staude, Navarro, et al., 2020; Staude, Waller, et al., 2020) will correlate with species declines in warmed treatments. We also predict that whether a species was already present at a destination site, potentially reflecting greater niche breadth and/or greater abundance, will be associated with more positive (or less negative) changes in abundance in warmed treatments (Lynn et al., 2021). These influences of rarity may also result in changes in phylogenetic diversity in turfs, if rarity is phylogenetically clustered. Previous work in this system found that species abundance showed phylogenetic signal (clustering) but range size did not (Veldhuisen et al., 2024), suggesting that initial abundance might be associated with phylogenetic consequences of community change, but range size influences on species’ responses to treatments are likely to be independent of phylogeny.

## Methods

### Study system

We conducted this experiment at three sites in Washington Gulch near the Rocky Mountain Biological Laboratory (RMBL, Gothic, Colorado, USA) from June to September 2017-2023. We do not have data from 2020, as the Covid-19 pandemic prohibited travel and fieldwork. RMBL is located in the East River valley of the West Elk mountains, approximately 10 kilometers from Crested Butte, Colorado. The relatively short growing season lasts 3-5 months, and snowmelt date is variable but has been arriving earlier on average with climate change (Aldridge et al., 2011; CaraDonna et al., 2014). Study sites were located at 2900 m (Low, 38.93248°N, –107.01018°W), 3150 m (Mid, 38.96046°N, –107.03262°W), and 3300 m (High, 38.97274°N, –107.04908°W) in elevation. The Low and Mid sites are in the montane vegetation zone, while the High is subalpine (Ackerfield, 2015). From the High to Low site, there is an increase in the growing season length, a mean growing season daytime air temperature difference of 1.9°C (2019 data from TOMST TMS-4 loggers placed at each site, Wild et al., 2019), and a mean growing season gravimetric water content difference of 0.17 g H2O per g dry soil. Fences were built around the sites to exclude cattle grazing.

### Experimental design & field data

In the fall of 2017, we transplanted turfs within and between each of the three sites (Fig. 1). We focused on grassland (grass/forb) vegetation and excluded shrubs when selecting turf locations. Each plot was 0.5 m x 0.5 m square and was moved intact to contain the entire plant community and an average of 20 cm of intact soil. To ensure replication, we divided sites into five blocks of five turfs each, arranged perpendicularly to the hillslope. In each block, we randomly assigned one of each treatment to each plot position (5 replicate turfs per treatment, 25 turfs total per site; Fig. 1). Treatments included: two ‘untouched controls’, a disturbance control that was locally transplanted, a one-or two-step warmer transplant (W1 and W2), and a one-or two-step cooler transplant (C1 and C2). “One-step” indicates that the turf was transplanted one site higher or lower, whereas “two-step” indicates the turf was transplanted two sites higher or lower. When transplanting, we maintained the orientation of each plot relative to the hillslope. Immediately after transplantation, we watered turfs from local streams and covered them with shade-cloth canopy for one month. In anticipation of natural disturbance from gophers, we added a sixth block of transplanted turfs (5 at the High site, 4 at the Mid site, and 6 at the Low site) in 2018, bringing the total to 90 turfs.

**Figure 1:**
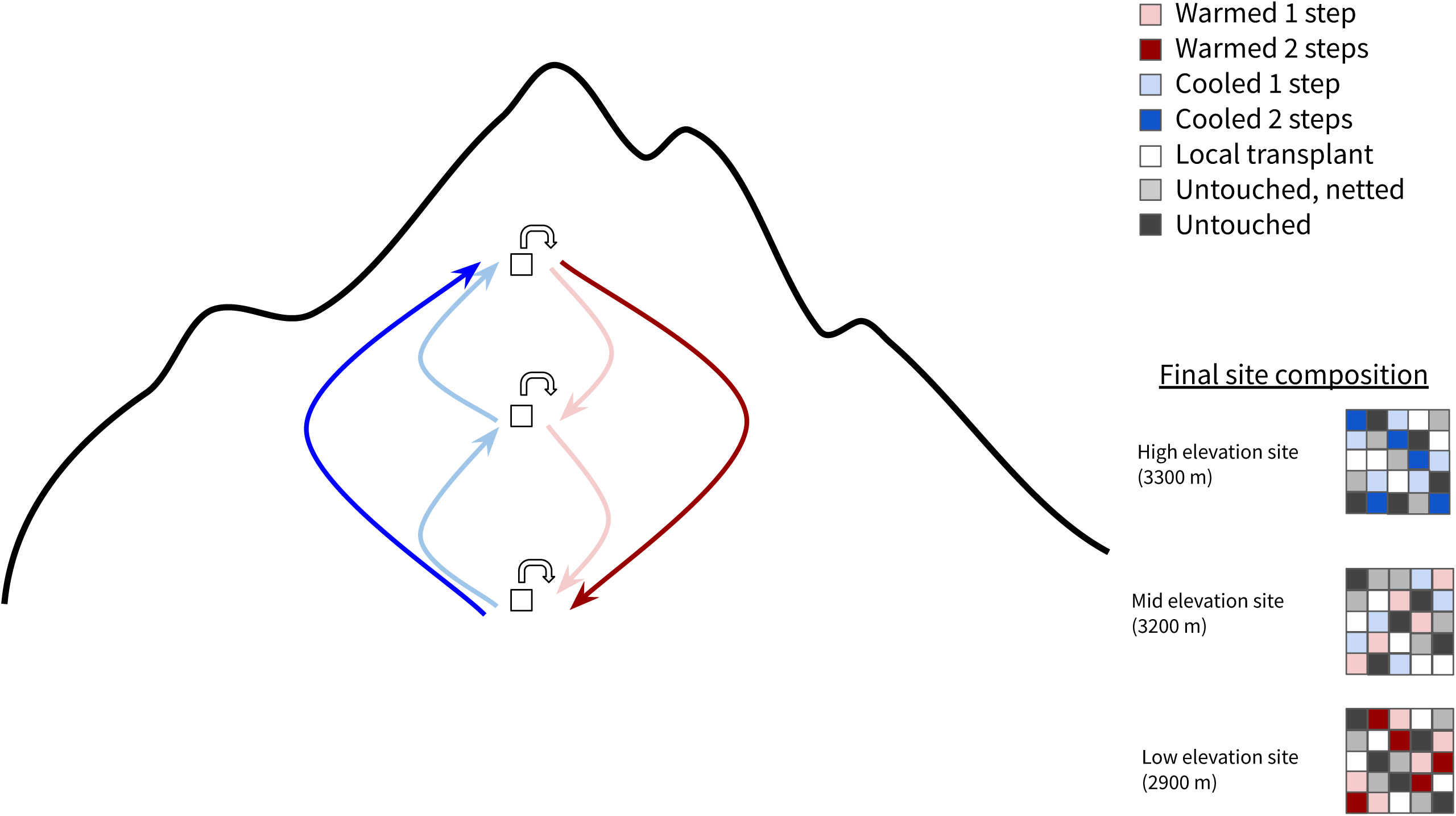
Experimental design. Each elevational site contains five blocks of five turfs transplanted between sites. Colors represent the treatment for each plot: warming (red), cooling (blue), or remaining at the same site as either fully untouched (dark grey), untouched, netted (light gray), or transplanted locally (white). Note: Colors are merely a depiction of the treatments in each block but do not accurately represent the location of each treatment within a block, which was randomized.

We installed nets around each turf (1 m x 1 m) in the growing season months to prevent gene flow from the transplanted plants to the surrounding plant communities. To preserve species interactions despite the nets, we centered each 0.25-square meter plot in a 1 m^2^ net, providing a buffer of destination site vegetation under the net. One untouched control was left un-netted in each block to test for the effect of nets. Analyses found no significant difference between the plant community variables of the three control types in this (L. Seltzer et al., in preparation, J. Navarro et al., in preparation) and similar turf transplant experiments(Henn et al., 2018), so here we exclude netted untouched and fully untouched control types, and only use local within site transplant controls for comparison to transplant treatments.

In 2017 only, before transplantation, we collected baseline plant species abundance data by counting individuals of each species in one turf (out of five) in each of the five blocks per site at time of peak plant biomass. From 2018-2023 (excluding 2020), we collected species percent cover data on all turfs for all species at the time of peak biomass in mid-late summer. We estimated the percent cover of each species in the plot to the nearest 1% using a 50 cm x 50 cm frame with a grid of 10 cm x 10 cm subsections. We also recorded species presence/absence in each subplot.

### Community-level metrics

We calculated community diversity metrics including species richness, Shannon diversity, and three phylogenetic diversity metrics for all turfs. We removed unknown species and did not include bare ground, rock, litter, or moss cover when calculating each metric. We focused only on those species who survived transplanting to the next year, and so we calculated diversity metrics for 2018-2023 (excluding 2020). To test prediction one, we calculated species richness by summing unique species in each plot with base R functions, and used percent cover data to calculate Shannon diversity in the R package ‘vegan’(Oksanen et al., 2024).

To quantify phylogenetic diversity to test prediction two, we calculated Faith’s phylogenetic diversity (Faith, 1992), mean phylogenetic distance (MPD) and mean nearest taxon distance (MNTD). We used the ‘ALLMB.tre’ angiosperm phylogeny from Smith & Brown (2018) to calculate all phylogenetic metrics. This phylogeny includes all genera in our dataset and 61 of 69 species observed in our turfs. Some species were only identified to genus during data collection, and we decided whether to drop them or substitute a different species in the phylogeny depending on whether congeners were in the turfs and if we could find genus-level phylogenies to inform substitutions (Appendix 1 Table S1). For species identified only to genus with no congeners in the turfs, we replaced them with a congener common in the area present in the Smith & Brown phylogeny. This substitution method applied to *Carex, Epilobium,* and *Lupinus*. Two genera (*Erigeron* and *Senecio*) had multiple species in the turfs, with some individuals only identified to genus and some species missing from the Smith & Brown (2018) phylogeny. *Senecio* had multiple species in the turfs missing from the phylogeny. We removed *Senecio crassulus* from all phylogenetic analyses because it has no available genetic or phylogenetic information. We substituted *Senecio serra* for the unidentified *Senecio* species since it is common in the area. We could not find an up-to-date genus-level phylogeny for *Erigeron* to inform replacements, so we removed *Erigeron coulteri* and the unidentified *Erigeron* sp. from all phylogenetic analyses. If a species was fully identified but missing from the phylogeny and was the only member of its genus in the turfs, we replaced it with a closely related congener common to the area in the Smith and Brown phylogeny (Appendix 1 Table S1).

We calculated Faith’s PD (Faith 1992), MPD, and MNTD in the R package ‘geiger’ (Pennell et al. 2014). Faith’s PD is the total evolutionary distance (branch length) across a phylogenetic tree describing the relationships among the taxa in a dataset. This metric will increase if relationships are more distant across the tree (species are more evolutionarily divergent). MPD is the mean distance between pairs of species, and so it also increases as evolutionary relationships are more distant across the tree, but will be less sensitive to rare cases of very distantly-related species that would otherwise elevate Faith’s Phylogenetic Diversity.

MNTD is the mean of the distance to the closest relative of each taxon, and so it reflects the typical distance to any species’ closest relative in the dataset and increases as closest relatives are more distant. Therefore, MNTD is more reflective of the tendency of species to have closer relatives than it is of the overall evolutionary diversity across the entire tree (community of species) that is captured by the former metrics (for further discussion, see review by Tucker et al. 2017).

To quantify whether a community had more phylogenetic diversity than expected by chance (overdispersion) or less (underdispersion), we calculated standard effect sizes (SES) for all three metrics in the ‘picante’ R package (Kembel et al. 2010) using the ses.metric() function. Positive SES values indicate that the community in question is more phylogenetically diverse than a random draw of the same number of species from the regional species pool (overdispersion), and *p* > 0.975 indicates statistical significance of the SES. Negative SES values indicate that the species group in question is less phylogenetically diverse than a random draw of the same number of species from the regional species pool (underdispersion), and *p* < 0.025 indicates statistical significance of the SES. *P* values are significant if the SES absolute value is greater than 1.96. We used each metric’s “sample.pool” null model to draw a random sample of the same number of species as the focal community from the pool of all species observed across all sites with 5000 iterations.

### Community-level analyses

To test the impacts of origin site, treatment and year on changes in the community diversity metrics to test predictions one and two, we used linear mixed models in the R package ‘lme4’(Bates et al., 2015). We used our plot-level calculations of species richness, Shannon diversity, PD, MPD and MNTD as response variables. We tested for individual effects of origin site, treatment, and year, interactive effects of treatment with origin site and year, and nested treatment within origin site. We then tested for significance in the interaction terms and compared AIC values to determine whether a nested model performed better than an additive model. We also set up a sum to zero contrast for origin site since no origin site is the obvious baseline comparison. We included plot number as random effect in all models to account for inherent variation between turfs. The final model we used for all metrics of community change is:

Community diversity _i_ = β_0_+ β_1_year + β_2_(origin site/treatment) + α_plot[i]_ + ε_i_; α_plot[i]_ ∼ N(0, σ^2^_α_))

where i denotes an individual plot, β values are the fixed effect coefficients, α is the random effect, ε_i_ is the random effect error term, and σ^2^ is the error variance.

### Species-level metrics

To quantify the change in individual species’ percent cover (Δcover_s_) in response to transplanting, we calculated the slope of the linear regression of cover versus year. We added zeroes for all years when a species was not observed in the turfs, and removed all species that were only observed in one year. We did not add zeroes for 2020, since no observations were made. We calculated average Δcover_s_ for each species in each treatment, and separated treatments if they were the same but their origin site was different. This resulted in a maximum of nine Δcover_s_ values for a species if it occurred in all three origin sites and turfs of all three treatments.

To quantify range size for each species, we calculated area of occupancy (AOO) with GBIF data. AOO correlates closely with population size and is thus considered more accurate than other range size metrics, and can be easily calculated from occurrence data (Gaston & Fuller, 2009; Staude, Waller, et al., 2020). To calculate AOO, we downloaded occurrence data for each species from GBIF using the ‘RGBIF’ package (Chamberlain & Boettiger, 2017) and calculated AOO with the ‘red’ package (Cardoso, 2017). We restricted GBIF data to occurrences in North America with <1000m coordinate uncertainty and observations that have been made since 1990 to ensure coordinate accuracy.

### Species-level analyses

To test prediction two, whether closely related species’ cover changed similarly in response to transplanting, we calculated phylogenetic signal in Δcover_s_ separately for each of the nine origin site/treatment groups. We used the ‘ALLMB.tre’ phylogeny from Smith & Brown (2018) and the same species substitutions as described in the “Community-level metrics” section above. We calculated phylogenetic signal for Δcover_s_, with Blomberg’s K (Blomberg et al., 2003) and Pagel’s λ (Pagel, 1999). We calculated Blomberg’s K with the “phylosignal()” function in the ‘picante’ R package (Kembel et al., 2010) with 5000 randomizations, and Pagel’s λ with the function “phylosig()” in the ‘phytools’ R package (Revell, 2011) with 5000 iterations and model set to “lambda.” Blomberg’s K or Pagel’s λ values near 1 indicate trait evolution consistent with Brownian motion. In contrast, K less than 1 or Pagel’s λ near zero suggests less phylogenetic signal than expected by Brownian motion.

To test prediction three, we tested whether species that originally occurred in their destination site prior to transplanting (those with higher habitat specificity) declined less than those that did not originally occur at their destination. For each of the nine origin site/treatment combinations, we grouped species by whether or not they occurred in their destination site pre-transplant according to the 2017 abundance data. We then used a t-test to compare the mean Δcover_s_ between the two groups.

To test whether a species’ pre-transplant (2017) abundance is associated with its Δcover_s_ to test prediction three, we ran a linear regression between the log of each species’ Δcover_s_ and its original abundance in that treatment group. We used the natural log + 1 of all 2017 abundances to avoid taking the log of zero. Finally, to test whether a species’ range size predicted its Δcover_s_, we conducted a linear regression of Δcover_s_ versus log of each species’ AOO.

All analyses were performed in R version 4.4.2 (R Core Team, 2024).

## Results

Across all sites and years, we recorded data for 69 species. Prior to transplanting, the Low elevation site had 55 species, the Middle had 52, and the High had 56, with significant overlap between all of the sites. By 2023, the last year of data collection, we recorded only 45 species across all sites and treatments, and 33 originally from the Low elevation site, 34 from the Middle, and 40 from the High. Despite the overall decline in richness across all sites and treatments, there were many species not recorded in certain sites pre-transplant that did occur in later years.

### Community-level change

We found no significant interaction between treatment and origin site or year in our linear mixed models, so all results here stem from models without interaction terms. We did find that the models with treatment nested within origin site performed better than additive models, so we report only results of nested models here.

Overall for prediction one, we found that species richness and Shannon diversity declined significantly in the cooled two steps treatment when compared with the local transplant turfs, while the warmed treatments generally increased in richness and Shannon diversity, but significance depended on the origin site (Fig. 2). Species richness declined significantly in turfs cooled one and two steps from the low elevation site (all *p* < 0.001), did not change significantly in turfs cooled one step from the middle elevation site (*p* = 0.41), and increased significantly in turfs warmed one and two steps from both the middle and high elevation sites (all *p* < 0.05; Fig. 2A). Shannon diversity declined significantly in turfs cooled one and two steps from the low elevation site (all *p* < 0.01), did not change in turfs cooled one from the middle elevation site (*p* = 0.91) or turfs warmed two steps from the high elevation site (*p* = 0.08), and increased significantly in turfs warmed one step from both origin sites (*p* < 0.05; Fig. 2B).

**Figure 2:**
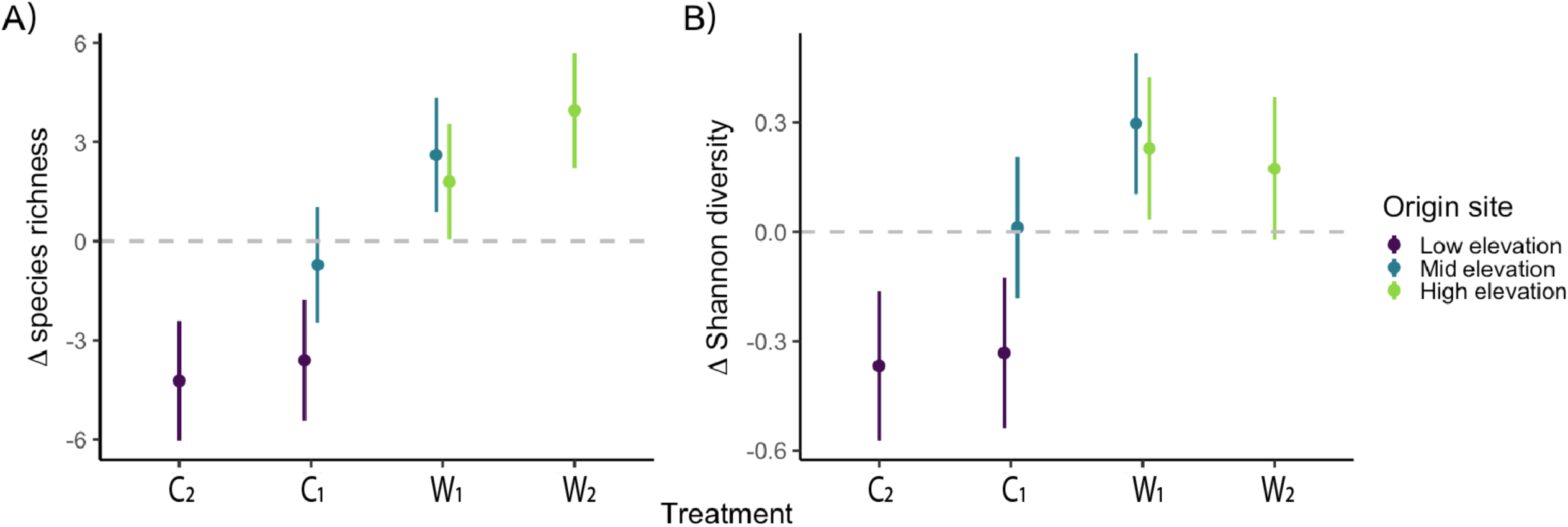
Changes in species richness. (A) and Shannon diversity (B) for each transplant treatment (where C is cooled, W is warmed, and subscripts denote 1 and 2 steps) relative to local transplant controls, whose change was set to zero (as indicated by grey dashed line). Bars are color coded by the origin site (Low site in purple, Mid in blue, and High in green). Points represent means ± SE.

For prediction two, the three phylogenetic diversity metrics exhibited largely opposite trends from richness and Shannon diversity, and generally increased in cooled treatments and didn’t change in warmed treatment (Fig. 3). PD increased significantly in all cooled turfs originating from the low elevation site (all *p* < 0.001), and increased but marginally significantly for both turfs cooled one step and warmed one step from the middle elevation site (C_1_ *p*=0.0533; W_1_ *p*=0.0496; Fig. 3A). PD did not change significantly for turfs warmed one or two steps from the high elevation site (all *p* > 0.05; Fig. 3C). MPD increased significantly in all cooled turfs regardless of origin site (all *p* < 0.05), and did not change significantly for any warmed plot from either origin site (all *p* > 0.05, Fig. 3B). MNTD increased in turfs cooled one and two steps from the low elevation site (all *p* < 0.001), as well as in turfs warmed one step from the middle elevation site (*p* = 0.01; Fig. 3C). MNTD did not change significantly in turfs cooled one step from the middle elevation site (*p* = 0.1) or in any warmed turfs from the high elevation site (all *p* > 0.05; Fig. 3C). Overall, turfs originating from the low elevation site (where only cooled treatments were possible) were the only ones that experienced significant differences in all five community diversity metrics regardless of treatment (Figs. 2 & 3).

**Figure 3:**
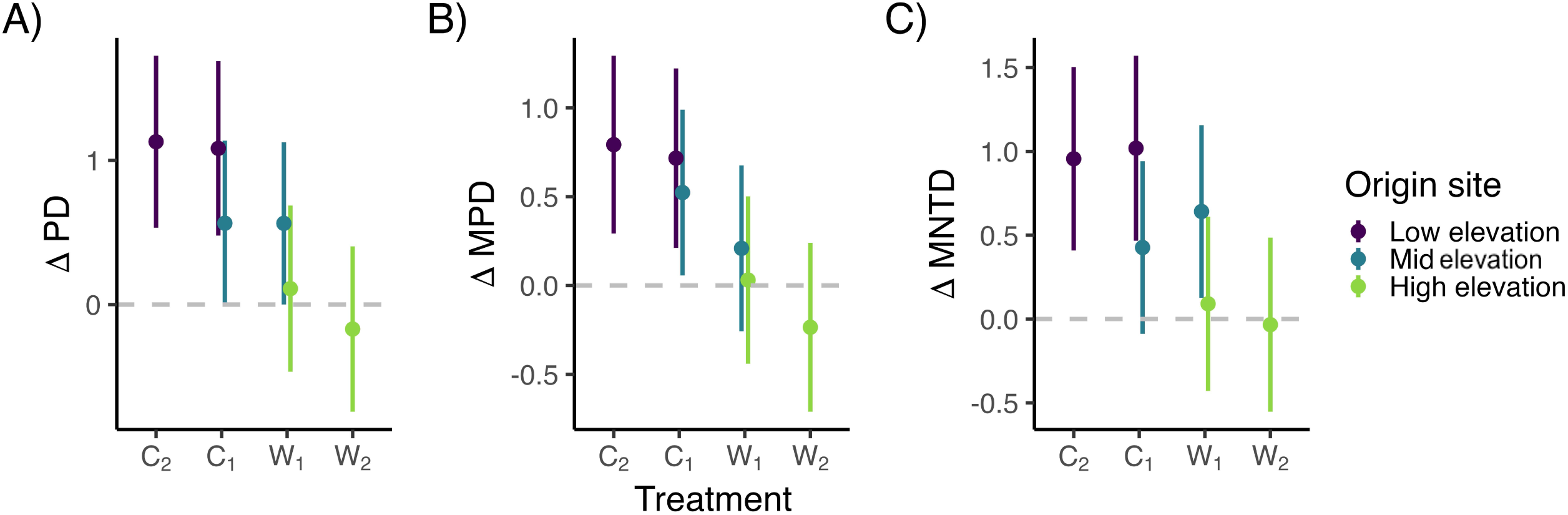
Changes in standard effect size for Faith’s phylogenetic diversity. (A), mean phylogenetic distance (B) and mean nearest taxon distance (C) for each transplant treatment (where C is cooled, W is warmed, and subscripts denote 1 and 2 steps) relative to local transplant controls, whose change was set to zero (as indicated by grey dashed line). Bars are color coded by the origin site (Low site in purple, Mid in blue and High in green). Points represent means ± SE.

### Species-level change

Across all origin sites and treatment groups, species’ Δcover_s_ from 2018-2023 ranged from –10 to 14, with a mean slope of 0.66 and median of 0.267 (Appendix 1 Fig. S1). We found no phylogenetic signal for Δcover_s_ in any of the origin site/treatment groups when looking at Blomberg’s K or Pagel’s λ (all *p* > 0.05; Fig. 4; Appendix 1 Table S2). For prediction three, we also found no significant differences in Δcover_s_ between whether species existed in their destination site pre-transplant or not (all *p*>0.05; Fig. 5). This pattern was the same for all origin site/treatment groups.

**Figure 4:**
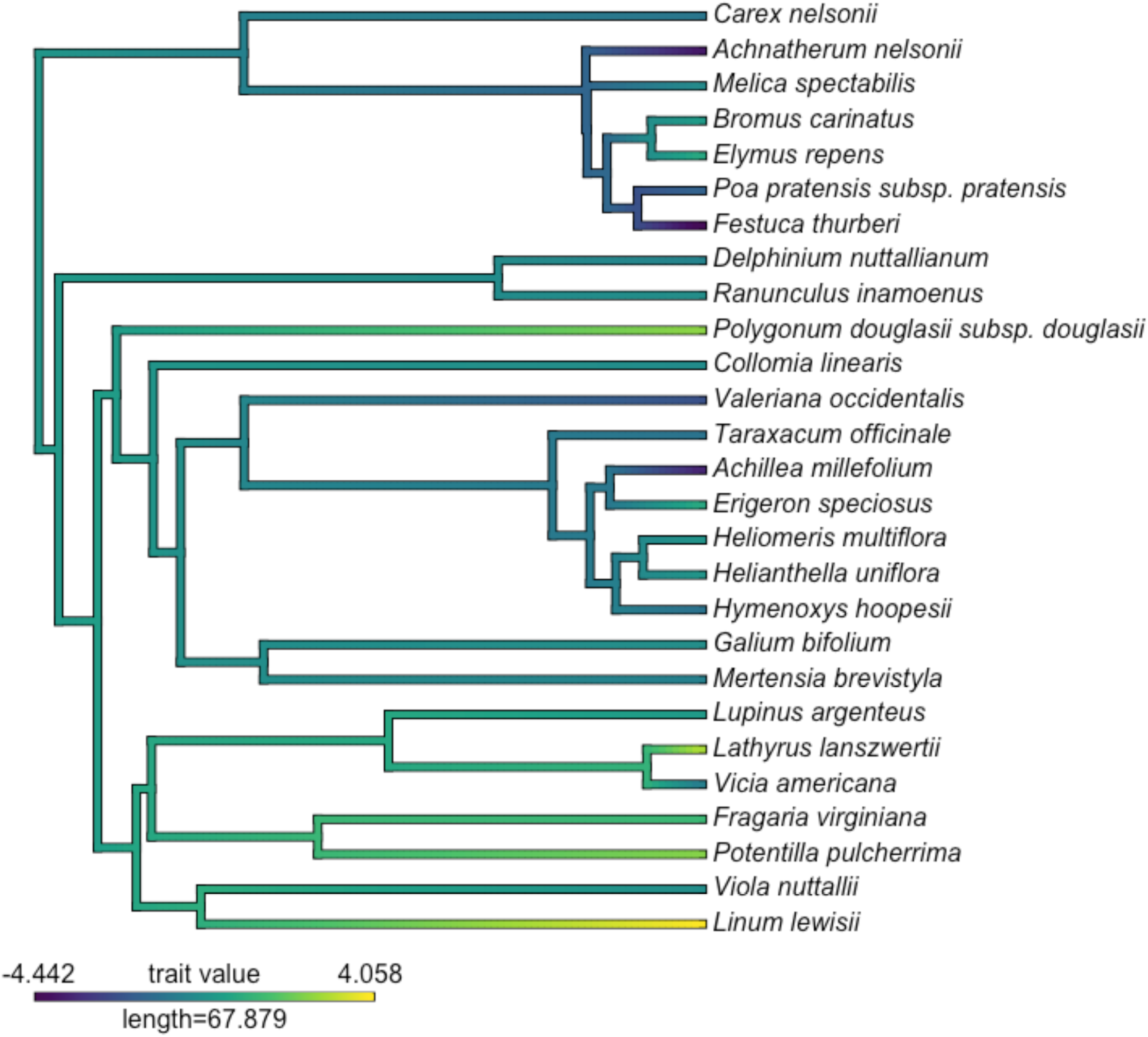
Phylogeny with the slope of species’ percent cover change over time indicated by branch color. This phylogeny shows phylogenetic signal for only the turfs cooled two steps at the Low site (see Appendix 1 Table S2 for all others; none show significant patterns). Cover change does not show any phylogenetic signal when measured by Blomberg’s K (K=0.326; *p*= 0.40) or Pagel’s λ (λ =0.210; *p*=0.365).

**Figure 5:**
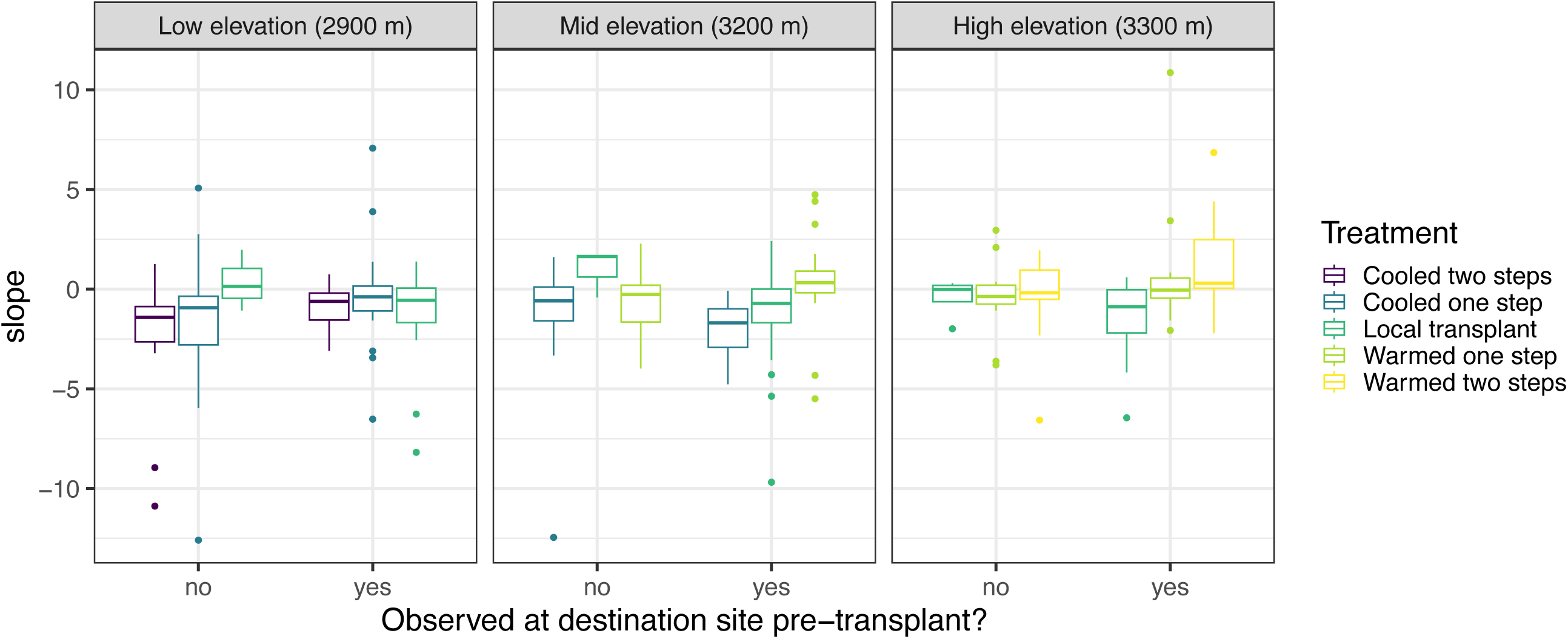
The slope of species’ cover changes, according to whether or not they occurred at their destination site pre-transplant. Panels are grouped by origin site (left to right: Low elevation to High elevation) and color coded by transplant treatment. There is no difference in species’ cover changes whether or not they occurred at their destination site pre-transplant for any origin site or treatment.

We found that 2017 pre-transplant abundance did not significantly predict Δcover_s_ for any of the treatment/origin site groups (all *p*>0.05; Fig. 6A). For the turfs warmed two steps from the high elevation site, we found a marginally significant, negative relationship between Δcover_s_ and 2017 pre-transplant abundance (R= –0.3; *p*=0.074; Fig. 6A). For all nine origin site/treatment groups, we found no significant relationship between Δcover_s_ and species’ range size (all *p* > 0.05; Fig. 6B).

**Figure 6:**
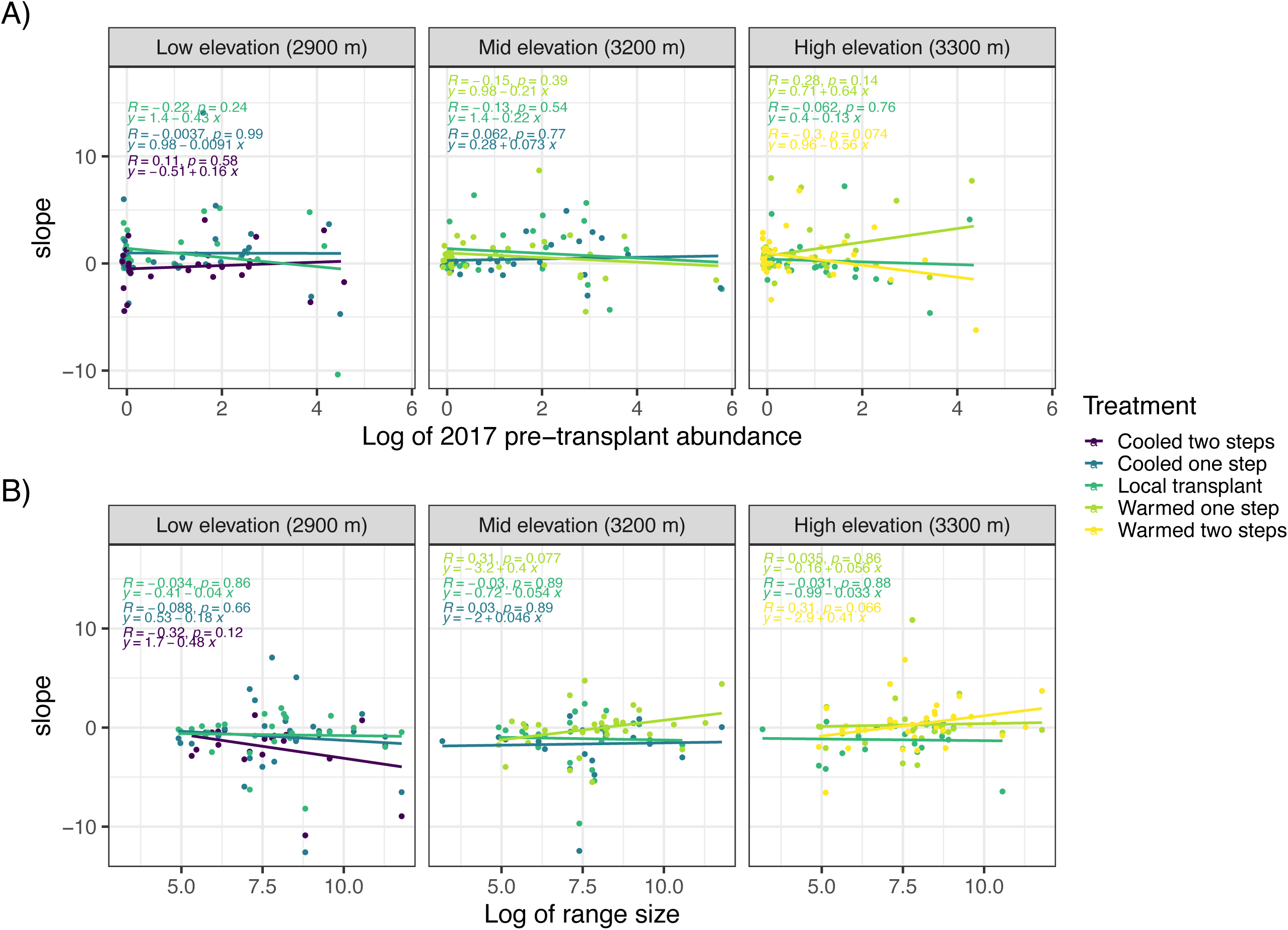
The slope of each species’ cover change versus its 2017 pre-transplant abundance. (A) and range size (B). Panels are divided by origin site from the Low site on the left to the High site on the right, and points are color coded by treatment. All relationships are nonsignificant. Points at x=0 are jittered slightly to make all visible.

## Discussion

How phylogenetic diversity responds to climate change in our Rocky Mountain plant communities, and to experimental transplanting globally, is largely unknown. We address this gap by testing how experimental warming and cooling via a whole community transplant experiment impacted species richness, Shannon diversity, and phylogenetic diversity. We predicted that 1) warming would drive declines in species richness and Shannon diversity and cooling would cause little change, 2) that warming would drive declines in phylogenetic diversity and cooling would again cause little change, and phylogenetic relationships would predict individual species’ responses, and 3) initial abundance, destination site occurrence and range size would also predict species’ responses to warming, with a potential association between phylogenetic relationships and abundance. Instead, we found that species richness and Shannon diversity largely increased with experimental warming and decreased with cooling, although significance depended on origin site. In contrast to richness and Shannon diversity, phylogenetic diversity by all metrics increased with cooling and didn’t change significantly with warming, though results again varied slightly by origin site. We found no significant phylogenetic signal in species’ cover changes for any treatment/origin site group. Finally, we found that species’ percent cover changes were not significantly predicted by pre-transplant occurrence at their destination site, pre-transplant abundance, or range size.

### Species richness and Shannon diversity increased under experimental warming, but not cooling

ur finding that species richness and Shannon diversity increased with warming was opposite of our prediction. Work tracking the impacts of both experimental and real climate change specifically in our Colorado Rocky Mountain system shows declines in diversity and abundance for many plant species (de Valpine & Harte, 2001; Iler et al., 2019; Panetta et al., 2018; Zorio et al., 2016), therefore we expected richness to decline in our warmed treatments. Work in other transplant systems has shown highly variable responses to experimental warming, including increases in colonization (Haider et al., 2022, 2024; Vandvik et al., 2020), increases in extinction and mortality (Bütof et al., 2012; Cui et al., 2018; Panetta et al., 2018; Walker et al., 2006; White et al., 2014), no significant changes in diversity metrics (Hudson & Henry, 2010; Shi et al., 2015), and responses that vary by location, species or functional group (Bjorkman et al., 2020; Elmendorf et al., 2012; Hudson & Henry, 2010; Vandvik et al., 2020; Wangchuk et al., 2021). Our results may align more with other transplant experiments around the world than with studies employing various methods in our same system.

Both increased colonization and lagged extinction, or ‘extinction debt,’ could contribute to increased richness. These effects have been observed in other systems (Alexander et al., 2018; Dullinger et al., 2012; Rumpf et al., 2019; Tilman et al., 1994), and have been documented in some transplant experiments (Haider et al., 2022, 2024; Vandvik et al., 2020). In a transplant experiment across 979 m in the German Alps, richness increased as warm-adapted species invaded warmed transplanted turfs, but extinction lagged (Haider et al., 2022, 2024). An initial study of this experiment spanned one year, while a follow up spanned four years (Haider et al., 2022, 2024). In the later study, the authors found that richness increased significantly at the beginning of their data collection, but those increases slowed over time (Haider et al., 2024). Similarly, our 5-year study duration may also have captured more colonization than extinction (Vandvik et al., 2020).

While both richness and Shannon diversity increased in all warmed turfs in our study, the increase in Shannon diversity was only significant in turfs warmed one step, and not those warmed two steps. The significant increase in richness in the turfs warmed two steps suggests that colonization is associated with a relative loss of evenness with greater warming, perhaps due to faster declines in abundance of species going extinct or slower increases in abundance among new colonists. Faster declines would be consistent with the more extreme treatment, and would predict that extinctions (and ultimately declines in richness) will be seen in these turfs first over longer periods of time.

### Community phylogenetic diversity increased under experimental cooling

In our turfs, phylogenetic diversity increased in cooled turfs and did not change significantly in warmed turfs, which was again contrary to our prediction of declines in warmed turfs and little change in cooled turfs. Additionally, our phylogenetic diversity changes were not in the same direction or magnitude as our species richness and Shannon diversity changes. There has been considerable debate about whether phylogenetic diversity is a reliable proxy for other biodiversity metrics (Cavender-Bares et al., 2004; E-Vojtkó et al., 2023; Gerhold et al., 2015; Hähn et al., 2024; Webb, 2000; Webb et al., 2002), and our results align with the more recent body of work suggesting that phylogenetic diversity cannot be assumed to parallel other diversity metrics and may respond differently to climate change (Daru et al., 2021; De Pauw et al., 2021; Hähn et al., 2024; López-Rubio et al., 2022).

Across all three diversity metrics, phylogenetic diversity largely increased in the cooled turfs, although the magnitude of the increase varied slightly by origin site. That cooling increased for all phylogenetic diversity metrics, while richness decreased, indicates that declining species are likely close relatives with species that are not declining, i.e. declining species are distributed across the phylogeny and are not clustered. The loss of close relatives would lead to increases in phylogenetic diversity overall, and particularly in the pairwise metrics, MPD and MNTD, as evolutionary distances between existing species increase. We also did not see declining phylogenetic diversity in warmed turfs, as would have been consistent with our prediction. The stable phylogenetic diversity with increasing richness and Shannon diversity indicates that newcomers are closely related to existing species. Overall, our results indicate that the identity of lost and gained species impacts phylogenetic diversity, and changes in individual species’ occurrences and species richness should be considered alongside phylogenetic diversity.

Previous work suggests that phylogenetic patterns could be explained by interactions with other species. The competition-relatedness hypothesis suggests that more closely related species compete more strongly (Darwin, 1859; Ricciardi & Mottiar, 2006). In line with this hypothesis, high phylogenetic diversity can be a signal of intense competition (Webb et al., 11/2002), although definitively linking phylogenetic diversity patterns with competition is complex, and studies have shown mixed support for this hypothesis (Bennett et al., 2013; Godoy et al., 2014; Lemos-Costa et al., 2024; Mayfield & Levine, 2010; Serván et al., 2024). Studies of transplant turfs have found that novel competitors strongly influence species’ responses to warming (Alexander et al., 2015; Nomoto et al., 2024; Nomoto & Alexander, 2021). In particular, Nomoto & Alexander (2021) found that novel competitors decreased warmed species’ population growth rates and reduced their predicted time to extinction, and that such competition may explain observed extinction debt. Notably, Nomoto and Alexander (2021) did not assess how relatedness may impact competition and population growth rate. Our results showed increasing phylogenetic distances between species in cooled turfs rather than warmed, which may not align with increasing competition in warmed turfs (Burns & Strauss, 2011; Gerhold et al., 2015; Mayfield & Levine, 2010; Webb et al., 2002). Additionally, we observed new species germinating post-transplant. Many species in these turfs reproduce clonally, and new individuals may have come from others just outside of the turf. New species could also have come from the pre-transplant seed bank, especially as the seasonal netting over our turfs would have somewhat limited local seed dispersal. These species that only germinated upon reaching their destination site may have created new competition dynamics. It would be valuable for future work in these turfs to quantify these interactions and link them with phylogenetic relatedness to understand the consequences for coexistence (Alexander & Levine, 2019; Van Dyke et al., 2022; Van Dyke & Kraft, 2025).

While we found many significant changes at the community level, we found no phylogenetic signal in individual species’ cover changes, indicating that species’ responses to warming and cooling via transplanting were not phylogenetically conserved. This result aligns with our finding of increased phylogenetic diversity in cooled turfs, again suggesting that species losses were distributed across the phylogeny, thus increasing relatedness between remaining species. To our knowledge, no other studies in transplant turfs have tested for phylogenetic signal in climate response, but other work has found that abundance change in response to 150 years of climate change was phylogenetically conserved in plants (Willis et al., 2008). Similarly, many studies have found evidence for phylogenetically-clustered extinction (Eiserhardt et al., 2015; González-Orozco et al., 2016; Willis et al., 2008; Zhao et al., 2024). This again suggests that additional time may be needed to observe abundance changes and their phylogenetic patterns. Continued testing for phylogenetic signal in experimental climate responses will be needed to assess whether closely related species face similar threats from ongoing global change.

### Aspects of rarity did not predict responses of individual species

Species’ cover responses were not associated with phylogeny, and they were also not correlated with their pre-transplant abundance or range size, or their pre-transplant presence in their destination site. These results are again inconsistent with our prediction. Low abundance is associated with extinction risk in many systems (Gaston & Fuller, 2008; Manes et al., 2021; Staude, Navarro, et al., 2020; Staude, Waller, et al., 2020), but it did not predict cover changes in transplant turfs in our study. None of the species in our turfs are listed as rare in the Colorado Rare Plant Guide (Colorado Rare Plant Guide, 2023), and as such, our range of species’ abundances may not have captured species with the most demographic sensitivity to changing conditions. Similarly, very few of our species had significant cover change at all. A previous study of this same system did find that local abundance has phylogenetic signal (i.e. abundance is clustered phylogenetically; (Veldhuisen et al., 2024), such that if initial abundance had predicted responses to treatments, cover changes would also have had a phylogenetic signal. Instead, cover changes did not have a phylogenetic signal, and phylogenetic diversity changes were consistent with phylogenetically distributed changes in species composition. These results are concordant with cover changes being decoupled from initial abundance.

Similarly, range size and presence at the destination site, indicators of a species’ niche breadth, did not predict their responses to transplanting across elevations. Range size is often expected to predict a species’ sensitivity to climate change (Pacifici et al., 2015; Schwartz et al., 2006). In Sonoran Desert plants, range size was not related to climate sensitivity(Brown et al., 2023), while in transplant turfs in Norway, it predicted colonization probability, but only in warmer and warmer/wetter turfs, not in only wetter or control turfs (Lynn et al., 2021). Previous work in Rocky Mountain plant communities suggests that range size does not show phylogenetic signal (Veldhuisen et al., 2024), so we expected that it may explain species’ phylogenetically dispersed declines. However, it did not. We also expected that whether a species existed already in its destination site would predict its cover change in response to transplanting, as local extinction risk was more closely correlated with climatic niche than range size in the Norwegian transplant turfs (Lynn et al., 2021). Lynn et al.’s (2021) work also showed that species with cooler niches were more likely to decline in warmed turfs, indicating that species whose destination site was already within their niche should do better than those whose destination site was outside of their niche. Oddly, our results were not consistent with this prediction, as we found no correlation between species’ cover change and existence at their destination site. This may be due to the length of our study, or the fact that many of our species occurred at multiple sites prior to transplanting, and thus reduced the power to detect an effect of presence at destination sites.

## Conclusions

Phylogenetic diversity is associated with ecosystem functioning, and is increasingly of interest as a target for conservation. Montane ecosystems experience intense impacts of climate change, but how these impacts affect phylogenetic diversity remains poorly understood. Our results suggest that species losses in the cooled turfs are phylogenetically dispersed, leading to increased phylogenetic diversity, even as richness and Shannon diversity decline. That phylogenetic diversity can increase with species loss is an important consideration for interpreting biodiversity impacts of community change. Meanwhile, increasing richness and Shannon diversity in our warmed turfs suggests that new species are able to colonize after the turfs have been transplanted, but also cautions that experimental turfs may also be experiencing extinction debt in their early years. Our results highlight that increased species richness and Shannon diversity do not translate to increased phylogenetic diversity, and vice versa, at least in the short term, emphasizing the importance of incorporating multiple biodiversity metrics into conservation decisions. Future work should address long-term changes in phylogenetic community composition, and recognize that the specific identities of lost and gained species will be essential to conserving phylogenetic diversity.

## Data Accessibility Statement

The data that support the findings of this study are available on Zenodo at https://doi.org/10.5281/zenodo.15643145. The Zendo reference number is 15643145. Code is available at https://anonymous.4open.science/r/transplant-plot-species-loss-263C/README.md.

