## Appendix 1 for "Phylogenetic diversity and species diversity are decoupled under experimental warming and cooling in Rocky Mountain plant communities"

*anonymous author names*

The data for these analyses is published on Zenodo: https://doi.org/10.5281/zenodo.15643145.

Table S1: Species in the plots listed with their replacements in the Smith and Brown (2018) ALLMB.tre phylogeny.

| Species in plots | in S&B? | Replacement | Notes |
|---|---|---|---|
| *Achillea millefolium* | yes | | |
| *Agoseris aurantiaca* | yes | | |
| *Agoseris glauca* | yes | *Agoseris glauca var. dasycephala* | subspecies |
| *Androsace septentrionalis* | yes | | |
| *Aphyllon fasciculatum* | yes | *Orobanche fasciculata* | synonym |
| *Aquilegia coerulea* | yes | | |
| *Artemisia tridentata* | yes | | |
| *Atriplex argentea* | yes | | |
| *Boechera stricta* | yes | | |
| *Bromus carinatus* | yes | | |
| *Campanula rotundifolia* | yes | | |
| *Carex sp.* | no | *Carex nelsonii* | common in area |
| *Castilleja sulphurea* | yes | | |
| *Chamaenerion angustifolium* | yes | *Chamerion angustifolium* | synonym |
| *Claytonia lanceolata* | yes | | |
| *Collomia linearis* | yes | | |
| *Delphinium nuttallianum* | yes | | |
| *Draba albertina* | yes | | |
| *Draba spectabilis* | yes | | |
| *Elymus bakeri* | yes | | |
| *Elymus repens* | yes | | |

| | | | |
|---|---|---|---|
| *Epilobium sp.* | no | *Epilobium ciliatum* | replacement, common in area |
| *Erigeron coulteri* | no | dropped | |
| *Erigeron elatior* | yes | *Erigeron grandiflorus* | synonym |
| *Erigeron sp.* | no | dropped | |
| *Erigeron speciosus* | yes | | |
| *Erythronium grandiflorum* | yes | | |
| *Festuca rubra* | yes | *Festuca rubra subsp. rubra* | correct subspecies |
| *Festuca thurberi* | yes | | |
| *Fragaria virginiana* | yes | | |
| *Frasera speciosa* | yes | | |
| *Galium bifolium* | yes | | |
| *Galium coloradoense* | yes | | |
| *Gentiana parryi* | yes | | |
| *Gentianella campestris* | yes | | |
| *Geranium richardsonii* | yes | | |
| *Helianthella quinquenervis* | no | *Helianthella uniflora* | replacement |
| *Heliomeris multiflora* | yes | | |
| *Heterotheca pumila* | no | *Heterotheca villosa* | replacement, common in area |
| *Hydrophyllum capitatum* | yes | *Hydrophyllum capitatum var. capitatum* | subspecies |
| *Hymenoxys hoopesii* | yes | | |
| *Lathyrus lanszwertii* | yes | | |
| *Ligusticum porteri* | yes | | |
| *Linum lewisii* | yes | | |
| *Lithophragma glabrum* | yes | | |
| *Lupinus sp.* | no | *Lupinus argenteus* | common in area |
| *Melica spectabilis* | yes | | |
| *Mertensia fusiformis* | yes | | |
| *Noccaea fendleri* | yes | | |

| | | | |
|---|---|---|---|
| *Osmorhiza occidentalis* | yes | | |
| *Poa pratensis* | yes | *Poa pratensis subsp. pratensis* | |
| *Poa reflexa* | yes | | |
| *Polygonum douglasii* | yes | *Polygonum douglasii subsp. douglasii* | subspecies |
| *Potentilla pulcherrima* | yes | | |
| *Ranunculus inamoenus* | yes | | |
| *Sedum integrifolium* | yes | *Rhodiola integrifolia* | synonym |
| *Senecio crassulus* | no | deleted | |
| *Senecio integerrimus* | yes | *Senecio integerrimus var. exaltatus* | subspecies |
| *Senecio sp.* | no | *Senecio serra* | replacement, common in area |
| *Stipa nelsonii* | yes | *Achnatherum nelsonii* | new taxonomy, new name |
| *Symphyotrichum ascendens* | no | *Symphyotrichum foliaceum* | replacement, common in area |
| *Taraxacum officinale* | yes | | |
| *Thalictrum fendleri* | yes | | |
| *Tragopogon dubius* | yes | | |
| *Valeriana capitata* | yes | | |
| *Valeriana edulis* | yes | | |
| *Valeriana occidentalis* | yes | | |
| *Veratrum californicum* | no | *Veratrum virginicum* | replacement |
| *Vicia americana* | yes | | |
| *Viola nuttallii* | yes | | |

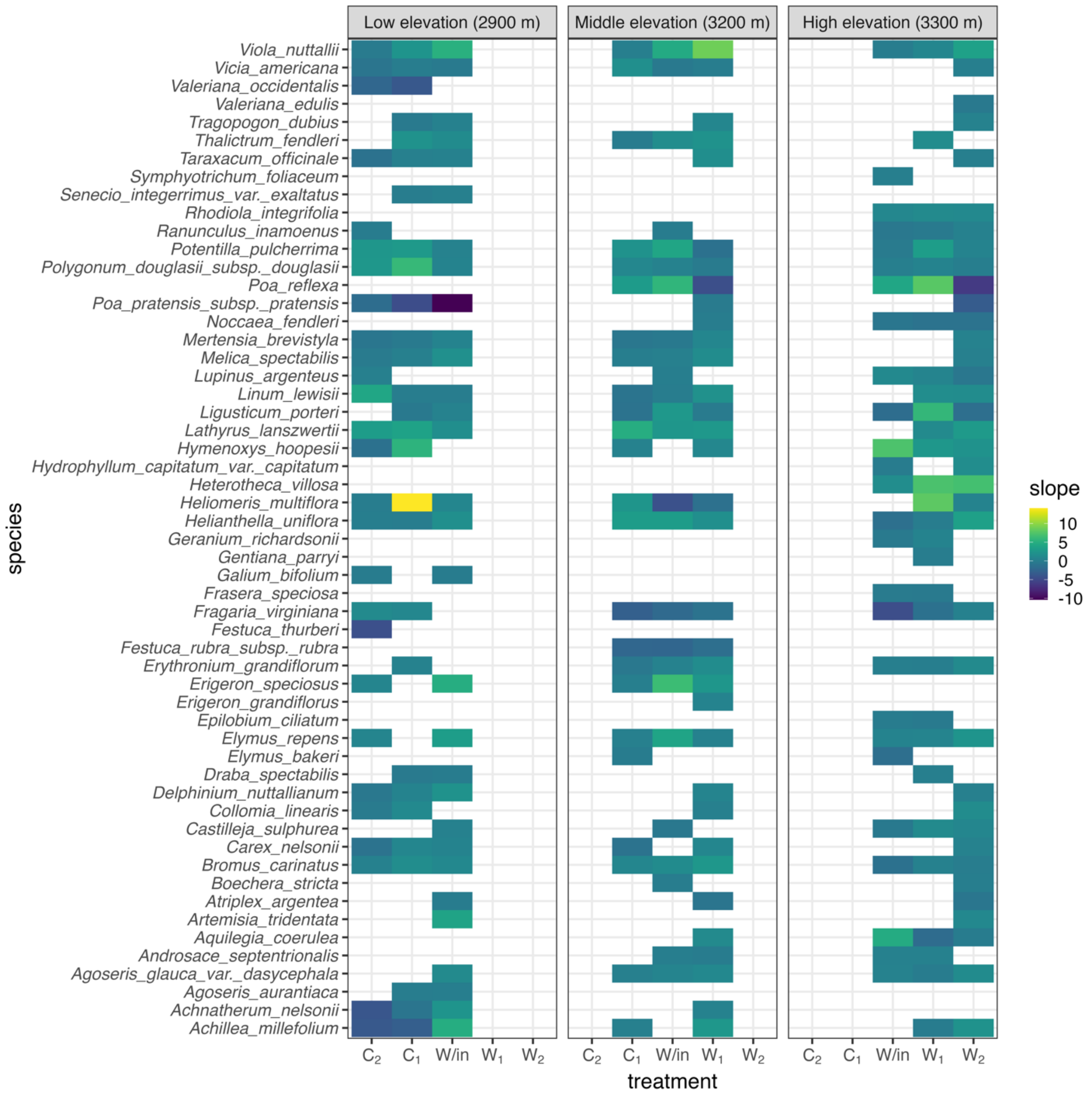


Figure S1: Heatmap faceted by origin site showing each showing the slope of each species' cover change over time (dark blue/purple for more negative changes to yellow for more positive changes).

Table S2: Phylogenetic signal values for species' percent cover change over time for all origin site and treatment combinations.

| Treatment | Origin site | Blomberg's K | P-value | Pagel's λ | P-value |
|---|---|---|---|---|---|
| Within site transplant | Low elevation | 0.22 | 0.55 | $7.3x10^{-5}$ | 1 |
| Cooled one | Low elevation | 0.19 | 0.94 | $7.3x10^{-5}$ | 1 |
| Cooled two | Low elevation | 0.33 | 0.4 | 0.21 | 0.36 |
| Within site transplant | Middle elevation | 0.17 | 0.99 | $7.3x10^{-5}$ | 1 |
| Cooled one | Middle elevation | 0.28 | 0.38 | $7.3x10^{-5}$ | 1 |
| Warmed one | Middle elevation | 0.12 | 0.6 | 0.009 | 0.96 |
| Within site transplant | High elevation | 0.22 | 0.64 | $7.3x10^{-5}$ | 1 |
| Warmed one | High elevation | 0.23 | 0.94 | $7.3x10^{-5}$ | 1 |
| Warmed two | High elevation | 0.24 | 0.56 | 0.14 | 0.244 |
